# Inverse mechanistic modeling of transdermal drug delivery for fast identification of optimal model parameters

**DOI:** 10.1101/2020.12.11.420836

**Authors:** Thijs Defraeye, Flora Bahrami, René M. Rossi

## Abstract

Transdermal drug delivery systems are a key technology to administer drugs with a high first-pass effect in a non-invasive and controlled way. Physics-based modeling and simulation are on their way to become a cornerstone in the engineering of these healthcare devices since it provides a unique complementarity to experimental data and insights. Simulations enable to virtually probe the drug transport inside the skin at each point in time and space. However, the tedious experimental or numerical determination of material properties currently forms a bottleneck in the modeling workflow. We show that multiparameter inverse modeling to determine the drug diffusion and partition coefficients is a fast and reliable alternative. We demonstrate this strategy for transdermal delivery of fentanyl. We found that inverse modeling reduced the normalized root mean square deviation of the measured drug uptake flux from 26 to 9%, when compared to the experimental measurement of all skin properties. We found that this improved agreement with experiments was only possible if the diffusion in the reservoir holding the drug was smaller than the experimentally-measured diffusion coefficients suggested. For indirect inverse modeling, which systematically explores the entire parametric space, 30 000 simulations were required. By relying on direct inverse modeling, we reduced the number of simulations to be performed to only 300, so a factor 100 difference. The modeling approach’s added value is that it can be calibrated once in-silico for all model parameters simultaneously by solely relying on a single measurement of the drug uptake flux evolution over time. We showed that this calibrated model could accurately be used to simulate transdermal patches with other drug doses. We showed that inverse modeling is a fast way to build up an accurate mechanistic model for drug delivery. This strategy opens the door to clinically-ready therapy that is tailored to patients.

## 1 INTRODUCTION

Mechanistic, physics-based modeling and simulation is increasingly used in healthcare for device engineering and design [1]–[3]. A typical example is the transdermal drug delivery systems (TDDS). Physics-based modeling was used to help design and optimize advanced TDDS with microneedles [4], [5], thermal skin ablation or iontophoresis [6], [7], chemical enhancers [8], and sonophoresis [9]. Also, first-generation TDDS are analyzed in-silico to increase our insight into how these systems interact with the patient’s skin [10]. Here, drug uptake differences between different substances [11] or age categories of patients [12] were targeted. Also, the impact of the composition of the stratum corneum and viable epidermis on the drug uptake was quantified [13], [14].

A key advantage of such first-principles-based modeling is the high spatial and temporal resolution they offer. Transient changes in concentration profiles can be identified within different skin layers during the uptake process [12], [15]. This spatiotemporal resolution enables us to virtually probe the drug transport inside and out of the skin in time and space. Also, the cumulative amount of drugs that are taken up by blood flow is available continuously, instead of only every 1-5 hours [12]. Another advantage is that such in-silico tests can be performed swiftly and noninvasively, without any side effects for the patient. Physics-based modeling opens new ways of tailoring TDD therapy for patients or patient groups [12]. This deterministic approach is also suffering from statistical variability in the data due to biological variability or measurement uncertainty. As such, only one in-silico experiment is required per case. Thereby, also very small differences between several cases can be accurately identified. Physics-based modeling and simulation provide valuable complementary information to experiments. Besides, they provide useful insights into the relative contribution of different drivers of drug uptake for the processes that can be captured mechanistically.

However, the reliability and accuracy of these physics-based models for TDD need to be verified and validated. FDA recently published guidelines for medical device design in this respect [1],[16]. A decisive aspect of determining model accuracy is the used model parameters. The properties of the skin, in particular, often play a crucial role, including drug diffusivities and partition coefficients [17].

There are three common ways of obtaining these properties of the skin. First, they are determined directly from exvivo measurements [18], [19]. Experimentally determining the diffusion and partition coefficients for a specific drug is resource- and time-consuming, especially when different skin types and anatomical locations on the human body need to be considered. Such an experimental campaign often poses a significant constraint on undertaking a modeling study. Thereby, they are rarely measured explicitly before modeling using a separate *in vitro* experiment. Second, skin transport properties for various molecules can be simulated with physics-based models [14], [20]. Here simulations at different scales are performed and coupled in a multiscale way [21], [22]. Even molecular dynamics simulations are applied [23]. Simulating, however, also consumes a lot of time and resources, making it is usually not more efficient than measuring the parameters separately. As a third way, modelers just source these parameters from literature data during model setup instead ([12], [24]–[26]). Due to a lack of appropriate data, often, data of other drugs with similar molecular weight and lipophilicity are taken [21]. Also, parameters are obtained from multiple sources. As these studies used different setups and boundary conditions (e.g., skin type, temperature, drug reservoirs), the absolute accuracy of the model results is often questioned. This is often the only way for modelers to simulate without a concurrent experimental or simulation campaign to determine material properties. The fourth way is using the quantitative structure-activity relationship (QSAR) modeling method to predict the biological activity of the new drug based on its structural properties [27]. This method can provide the stimation of different values for properties of subtances that were not studied experimentally [28], however, the predicted value is limited to the constraints of the implemented dataset [29].

There is a fifth alternative currently underexplored. All skin material properties could be fitted simultaneously to obtain the best agreement with measured data of the drug uptake kinetics. Calibrating a mathematical model by comparing its predicted response (i.e., drug uptake) with experimental observations is called inverse modeling [30]. This strategy ensures that the in-silico system is calibrated so that it responds as close as possible in the same way as the in vitro (or in vivo) system. A key advantage is that only limited experiments on the drug uptake kinetics need to be performed, instead of performing separate experiments for each property on a different apparatus. A step in this direction was made [31]. The diffusion and partition coefficient for the transport of flufenamic acid through the stratum corneum layer were determined by fitting the simulations to the experiments. The *in vitro* concentration-depth profiles were used to calibrate these material properties. A nonlinear least-squares approach was used to determine these parameters both for every point in time or averaged over all measured times. However, this study did not target the drug uptake amount through the skin by the patient and did not include the transport properties of the patch as possible unknowns that could be fitted.

This study aims at developing and testing a fast and straightforward way to calibrate a physics-based model for transdermal fentanyl delivery. Fentanyl patches for around-the-clock opioid analgesia are currently amongst the most popular transdermal delivery devices [32]. We target to have an excellent agreement of our model for drug uptake kinetics using a commercial fentanyl patch currently used in the clinics. To this end, a sizeable parametric space of the skin epidermis’ material properties and the patch is simulated for transdermal fentanyl uptake through a Franz diffusion cell. A comparison with experimental data on the drug flux out of the epidermis is made. This enables to elucidate *in silico* the sensitivity of the drug uptake kinetics to each model parameter in the transdermal patch and the skin. Within this parametric space, we determine the best set of properties with a better agreement with experiments during a 72-h drug uptake period, compared to relying on experimentally-determined skin properties. A faster alternative procedure to multiparameter inverse modeling by the parametric exploration is also explored, using automated least-squares optimization. The validity of the obtained parameter set for simulating other drug patches with a different concentration is also tested. This independent verification proves the generalized use of the model.

## 2 MATERIALS AND METHODS

### 2.1 Continuum model for transdermal fentanyl delivery

An extensive mechanistic model was set up to simulate fentanyl release from a transdermal patch (reservoir) and subsequent uptake through the human epidermis [12]. The model and the corresponding simulation were built and executed according to best practice guidelines in modeling, among others, those for medical device design [1], [33]. The model was used to simulate the drug uptake experiment described in [18]. By comparison with the experimental data, we can quantify how accurately the drug uptake kinetics are predicted with our mechanistic model.

#### 2.1.1 Experimental setup and data

The experiment involved fentanyl uptake from a cylindrical drug reservoir (radius 9 mm, thickness 50.8 μm) through a cylindrical skin sample (human cadaver epidermis, viable epidermis, and stratum corneum, radius 9.25 mm, thickness 50.8 μm, Figure 1a) into an aqueous solution. The drug was embedded in an acrylate polymer, which served as the donor reservoir. Both the reservoir and epidermis were fitted into a Franz diffusion cell and kept at 33°C. The receptor medium in the cell was a phosphate buffer at a pH of 5.65, in which fentanyl had a solubility of 2.5 mg ml^−1^. The experiments took 72 h. Several initial drug concentrations in the patch (*c_pt,ini_^α^*) were evaluated for fentanyl (substance *α*). We used the data on the drug uptake fluxes that left the epidermis for *c_pt,ini_^α^* = 60 kg m^−3^and *c_pt,ini_^α^* = 80 kg m^−3^. This flux in the experiments was determined at discrete points in time. An aliquot of the receptor medium was removed for this purpose. Its concentration was analyzed via high-performance liquid chromatography (HPLC). As such, the average flux over a specified time period of multiple hours was obtained.

**Figure 1.**
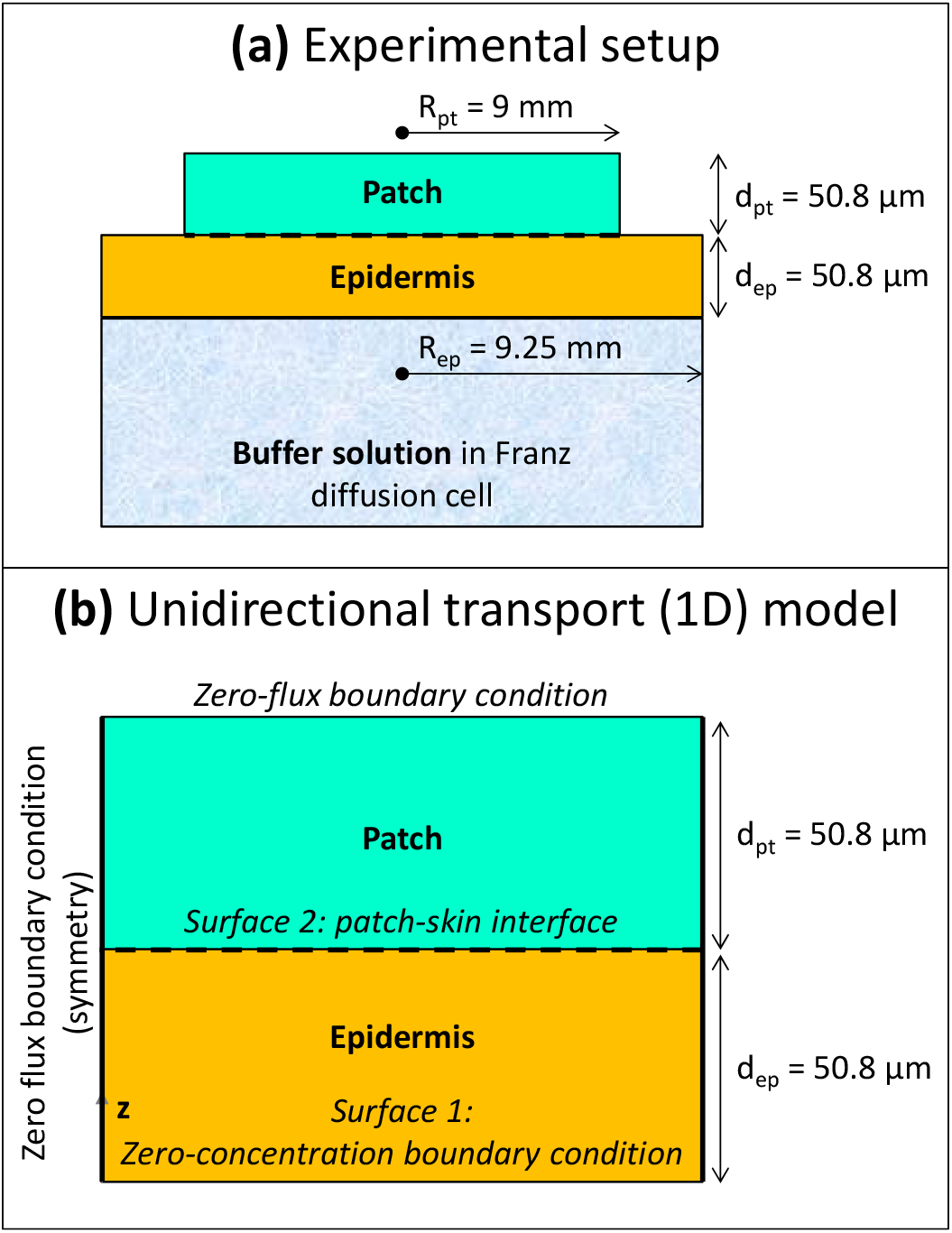
The geometry of the experimental setup (a) and of the 1D mechanistic model (b) of a cylindrical drug reservoir and epidermis (not to scale).

#### 2.1.2 Computational system configuration

The geometrical setup is depicted in Figure 1b, along with the boundary conditions. The system configuration is built up, similar to the experiment [18]. The configuration includes a cylindrical drug reservoir and the outer part of the human skin, namely the epidermis. The reservoir contains a finite amount of fentanyl. The geometrical specifications and transport properties used in the simulations are indicated in Table 1.

**Table 1.**
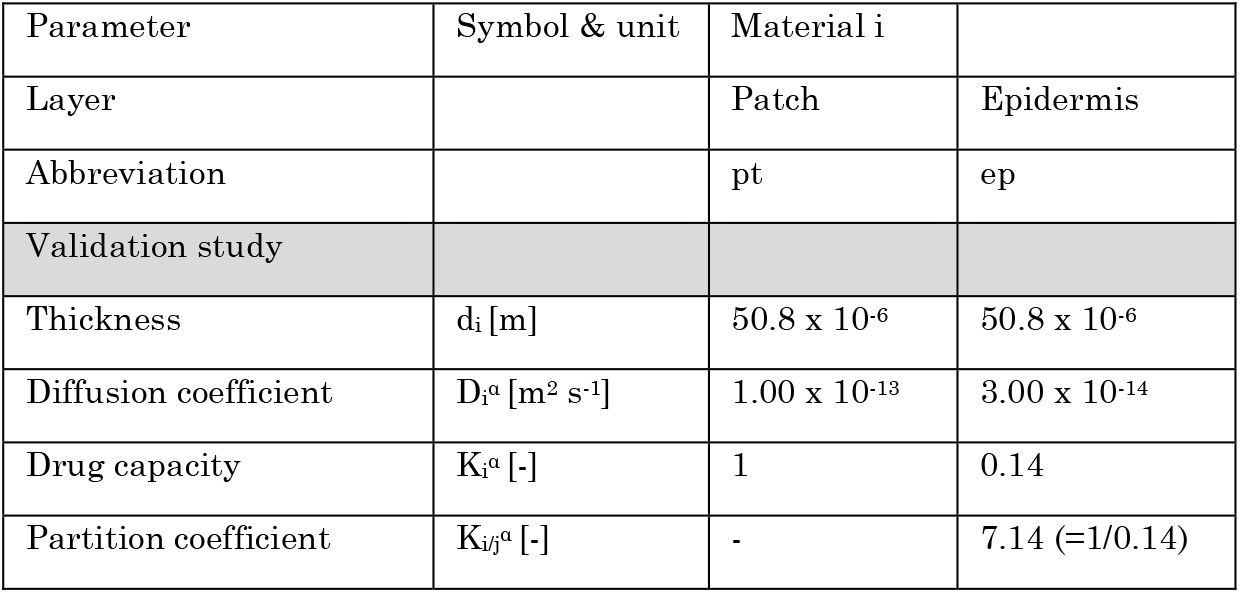
***Material transport properties are used in the model for the base case for fentanyl from*** [18].

| Parameter | Symbol & unit | Material i |  |
| --- | --- | --- | --- |
| Layer |  | Patch | Epidermis |
| Abbreviation |  | pt | ep |
| Validation study |  |  |  |
| Thickness | $d_i$ [m] | $50.8 \times 10^{-6}$ | $50.8 \times 10^{-6}$ |
| Diffusion coefficient | $D_i^a$ [$m^2 s^{-1}$] | $1.00 \times 10^{-13}$ | $3.00 \times 10^{-14}$ |
| Drug capacity | $K_i^a$ [-] | 1 | 0.14 |
| Partition coefficient | $K_{ij}^a$ [-] | - | 7.14 (=1/0.14) |

Given the very similar radius of the patch and the skin sample in the experiment, the transport processes can be considered one-dimensional. This led to minimal differences when comparing the 3D and 1D model representation [12]. Differences in predicted fluxes and the cumulative drug amount that was taken up by the aqueous buffer solution were < 0.4%. Thus, a 1D model was sufficiently accurate and improved the computational economy, which was beneficial because a large parametric space was explored. Such a 1D model is representative of Franz diffusion cell experiments [18], [34], [35]. The skin is as wide as the drug reservoir, so unidirectional (longitudinal) drug transport is simulated without any transverse component.

A patch radius Rpt of 9 mm was used for the drug reservoir, similar to in the experiments. This value implies an active area of 2.54 cm^2^, which is in the same order of magnitude as reported for commercial transdermal patches for fentanyl (4.2-42 cm^2^) [32]. The thickness of the patch (*d_pt_*) was 50.8 μm. The volume of the patch reservoir was 1.29 x 10^−8^ m^3^. The skin’s epidermis (ep) was modeled as a single layer. This approach lumps the markedly different diffusion and partitioning processes through the lipophilic stratum corneum and the hydrophilic viable epidermis. This strategy does not affect the uptake kinetics for such a one-dimensional setup without extensive lateral transport. As such, a realistic simulated uptake drug flux across the skin into the Franz diffusion cell g_bl,up_(t) [kg m^−2^ s^−1^] is obtained. It is the performance metric used to compare with experimental data. However, this one-layer approach does not accurately reflect the spatial concentration gradients within the skin, as shown in [12], where much higher concentrations are found within the stratum corneum.

#### 2.1.3 Governing equations

The governing equations for simulating transient transport of fentanyl are detailed. Only drug diffusion was solved, and isothermal conditions were assumed, close to human body temperature. Water transport due to skin de- /rehydration and the resulting skin shrinkage or swelling were not modeled. The following mass conservation equation was derived for fentanyl (substance *α*) in [12]. This equation is defined for each material *i* [kg m^−3^] (patch and epidermis) to calculate the drug potential *ψ^α^*:

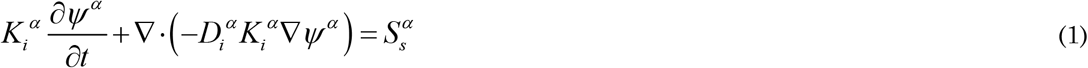

*D_i_^α^* is the diffusion coefficient or diffusivity [m^2^ s^−1^], *K_i_^α^* is the drug capacity of the drug in the material *i* [-], *S_s_^α^* is a volumetric source term for substance α [kg m^−3^s^−1^], and *t* is the time [s].

The conservation equation was derived in drug potential (*ψ^α^*) instead of drug concentration (*c_i_^α^* = *K_i_^α^ ψ^α^*). The use of *ψ^α^* instead of *c_i_^α^* leads to a single dependent variable throughout the entire domain, instead of one in each material. This choice mainly avoids numerical stability issues at the interface of the patch and epidermis [12]. Such instabilities can occur due to the discontinuity in the drug concentration that arises at the interface caused by partitioning.

Partitioning implies that when a drug substance α is brought into contact with patch and epidermis (pt and ep), the drug will evolve to a different equilibrium concentration in each of these materials, namely *c_pt_^α^* and *c_ep_^α^*. The ratio of these equilibrium concentrations is referred to as the partition coefficient *K_pt/ep_^α^*.

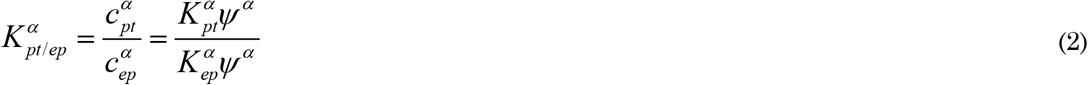

The octanol-water partition coefficient is frequently determined (*K_o/w_^α^*) for drug partitioning in liquids, where values larger than 1 indicate drug lipophilicity. Values smaller than one indicate drug hydrophilicity. The log(*K_o/w_^α^*) is often reported, where positive/negative values indicate lipophilicity/hydrophilicity, respectively.

No source term was included in this study. The reason is that the contributions of the following processes [10] could be neglected for fentanyl, as motivated in [12]: (1) metabolization of the drug molecule within the epidermis by a chemical reaction, a process that leads to a conversion of the drug into other compounds; (2) adsorption of the drug molecule into the epidermis, and thus physical binding of the drug molecules.

#### 2.1.4 Material properties and transport characteristics of skin and patch

Fentanyl is a synthetic opioid that is used as a pain medication. It has a low molecular weight (337 Da) and is moderately lipophilic with a log(*K_o/w_*) of 3 to 4 [21], [32], [36]. The material transport properties of the skin components and the drug reservoir are given in Table 1 for fentanyl for the base case, as taken from [18]. These parameters were estimated in [18] as follows:

- The diffusion coefficient of fentanyl in the patch (*D_pt_^α^*) was determined experimentally. This was done by performing a drug release experiment of the patch into a receptor medium, after which an analytical expression was fitted to the results.
- The drug capacity of fentanyl in the epidermis (K_ep_^α^ = 0.14) was derived from sorption experiments, measured over five skin donors. The patch’s drug capacity (K_pt_^α^) was taken equal to one since it is taken as the reference domain. Note that in [18], these drug capacities are actually termed partition coefficients, which is not the same definition used in the present study. The actual partition coefficient between patch and epidermis K_pt/ep_^α^ = K_pt_^α^/K_ep_^α^ equals 1/0.14.
- The value for the baseline diffusivity of the drug in the epidermis (D_ep_^α^) was determined by fitting the experimentally measured drug flux in the Franz diffusion cell over a range of initial drug concentrations in the patch for the initial 30 h [18].

These different procedures illustrate the complexity of obtaining material transport properties for transdermal drug delivery. Three different experimental setups were required, where even model fitting was required for one parameter (D_ep_^α^). In addition, each experimental technique had specific uncertainty.

Similar to most other simulation studies [15], [21], [22], the diffusion and partition coefficients were taken as constants and isotropic. This assumption implies that they are independent of the drug concentration. This choice is often unavoidable because more detailed data is rarely available. However, diffusion and partition coefficients have been shown to be a function of drug concentration rather than constant values [23].

#### 2.1.5 Boundary and initial conditions

The drug was assumed to exit the computational domain via the interface with the aqueous solution in the Franz diffusion cell. Therefore, a constant concentration (and potential), equal to zero, was set at the epidermis bottom (Figure 1). This approximates the very low concentration found in the large buffer solution. This condition represents a Dirichlet boundary condition. Zero-flux conditions were imposed at all vertical boundaries. At t = 0 s, the skin was assumed to be drug-free. The initial concentration of drugs in the patch was set at 80 kg m^−3^ or 60 kg m^−3^, according to a previous study [18]. The patch dimensions combined with these initial concentrations lead to an initial amount of fentanyl in the reservoir (m_pt,ini_) of 1.03 mg and 0.776 mg, respectively. This initial drug content corresponds to amounts typically present in commercially available patches [12]. Perfect contact between the patch and the epidermis was assumed. No inclusion of air layers or discontinuities like hairs or skin roughness were accounted for.

### 2.2 Spatial and temporal discretization

The grid was built based on a grid sensitivity analysis. The spatial discretization error was quantified based on the total mass flux to the Franz diffusion cell. This error was estimated to be 0.1%, as determined by Richardson extrapolation [37], [38]. The grid consisted of 112 quadrilateral finite elements for the base case (1D, elements with a size of about 1 μm). The grid was gradually refined towards the patch-skin interface to enhance numerical accuracy and stability. The reason is that the largest gradients are found at such interfaces, particularly at the uptake process initial stage.

The transient simulations quantified a drug uptake process that lasted 72 h (3 days), starting from these initial conditions. This time window also agrees with that of conventional transdermal fentanyl therapy. Here the required dose for the patient, so patch size is estimated empirically by the clinician. Afterward, the patch is applied transdermally and is replaced every 72 hours [39].

The simulations applied adaptive time-stepping- The maximal time step was 600 s (10 min). This time step ensured high temporal resolution for the output data and was determined from a sensitivity analysis.

### 2.3 Numerical implementation and simulation

The model was implemented in COMSOL Multiphysics^®^ software (version 5.5, COMSOL AB, Stockholm, Sweden). COMSOL is commercial finite-element-based software. This software was verified by the code developers. Therefore, additional code verification was not performed by the authors. Transient diffusive drug transport (Eq.(1)) in the patch and epidermis was solved using the partial differential equations interface (coefficient form). The conservation equation was solved for the dependent variable *ψ*. The optimization interface was used to automatically find the optimal model-parameter values by direct inverse modeling [40]. Here, the least-squares method was used. This method minimizes the objective function, namely, the root mean square of the differences between experimental data and simulations. Different starting points were applied to avoid landing in a local minimum. In addition, the results were compared to the results of the full parametric space as well to ensure no local minimum was identified.

Quadratic Lagrange elements were used together with a fully-coupled direct solver, which relied on the MUMPS solver scheme (MUltifrontal Massively Parallel sparse direct Solver). For the direct inverse modeling, a derivative-free solver was used, namely the Bound Optimization by Quadratic Approximation (BOBYQA). The tolerances for solver settings and convergence were determined through sensitivity analysis so that a further increase in the tolerances did not alter the resulting solution. For the inverse modeling, different starting values of the material properties were explored, and the sensitivity to the stopping criterion parameters was also performed.

### 2.4 Parametric study

Following simulations were performed:

1. The base case simulates the drug uptake through the epidermis as released from a finite drug reservoir based on experimentally determined material properties (Table 1) [18]. Simulations were done for c_pt,ini_^α^ = 80 kg m^−3^.
2. Indirect multiparameter inverse modeling to determine the optimal set of material properties (D_pt_^α^, D_ep_^α^, K_ep_^α^) that gives the best agreement with the experimental data during a 72-h drug uptake period (section 3.1) for c_pt,ini_^α^ = 80 kg m^−3^. Indirect modeling implies that a large set of combinations of these properties was explored; in this case, 30 618 combinations/simulations. The main aim here was to get insight into how the relationship between these parameters affects solution accuracy.
3. Direct multiparameter inverse modeling to determine the optimal set of material properties (D_pt_^α^, D_ep_^α^, K_ep_^α^) that gives the best agreement with the experimental data (section 3.2) for c_pt,ini_^α^ = 80 kg m^−3^. Here automated least-squares optimization was used to progress fast to the most optimal solution. The main aim here is to identify how much faster a solution can be found compared to indirect inverse modeling. We also performed direct inverse modeling for c_pt,ini_^α^ = 60 kg m^−3^.
4. Cross verification of the obtained parameter set by direct inverse modeling (for c_pt,ini_^α^ = 80 kg m^−3^) by comparison with independent experimental data of patches with a different drug concentration (c_pt,ini_^α^ = 60 kg m^−3^). This independent verification proves the generalized use of the model, so if the calibrated model can be used for a wider operational range of patches. We also performed the same cross-verification by evaluating a patch with an initial drug concentration of c_pt,ini_^α^ = 80 kg m^−3^ using the optimal model parameters obtained from direct inverse modeling for c_pt,ini_^α^ = 60 kg m^−3^.

### 2.5 Metrics to evaluate TDD

The simulated drug delivery process was analyzed quantitatively by calculating several metrics. From the experiments, the flux taken up by the Franz diffusion cell (g_Fr,up_ [kg m^−2^ s^−1^]) was measured as a function of time for discrete points in time. This experiment was done by measuring the total amount of drugs taken up in the respective timeframe into the Franz diffusion cell through surface 1 [kg s^−1^]. Afterward, this flow rate was scaled with the patch’s surface area (surface 2, *R_pt_* = 9 mm). This flux quantity was used as the primary metric to compare with the 1D simulations, where it was determined in the same way. The Root Mean Square Deviation (RMSD) between measured fluxes and the predicted fluxes by the simulations over the N measured points at each time *j* was calculated as:

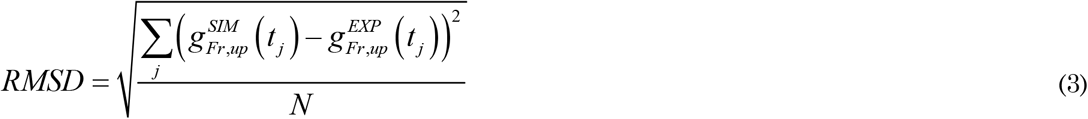

This RMSD [kg m^−2^ s^−1^] is the quadratic mean over these differences in uptake fluxes between experiments and simulations. Thereby it is a direct measure of the accuracy of the simulations. Using the RMSD, we can quickly elucidate how far off these parameters are from the experimental data and the original dataset (base case). The advantage of the RMSD is that it aggregates each simulation’s accuracy into one single value, being the prediction error of the simulation. An additional benefit of the RMSD for comparison is that it has the same units as the flux [kg m^−2^ s^−1^], so it can be directly contrasted to the measured fluxes. The RMSD was calculated considering the entire 72-hour timeframe (N=9 data points) and a shorter 30-hour timeframe (RMSD’, N=4).

In addition to the RMSD, the average uptake flux across the skin into the diffusion cell over the 72 hours is quantified 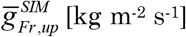. The reason is that the normalized RMSD (NRMSD) can be calculated as the ratio of the RMSD to this average flux, which can be expressed as a percentage as well:

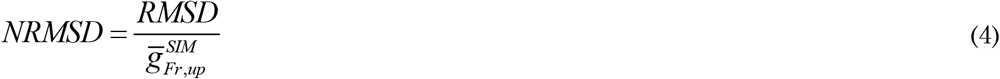

## 3 RESULTS AND DISCUSSION

### 3.1 Sensitivity of drug uptake kinetics to material properties

We aim to quantify how the model parameters – the material properties D_pt_^α^, D_ep_^α,^ and K_ep_^α^ – affect the simulations’ accuracy, so the RMSD. We mainly aim to elucidate if there are particular combinations of model parameters that enhance model accuracy. These results shed light on which transport kinetics are decisive for achieving an accurate mechanistic model. Different subsets of the material properties were analyzed within the sizeable parametric space explored for D_pt_^α^, D_ep_^α,^ and K_ep_^α^. First, we analyze the impact of epidermal diffusion versus partitioning in the epidermis. Second, the effect of diffusion in the epidermis versus diffusion in the patch is targeted. Finally, the most accurate combination of parameters in this design space that leads to the lowest RMSD is identified.

#### 3.1.1 The impact of epidermal diffusion and partitioning

Parameter variations in the diffusion (D_ep_^α^) and partition coefficient of the epidermis (K_ep_^α^) are explored. The diffusion coefficient in the patch (D_pt_^α^) is kept the same as the base case. The resulting RMSD values for each combination of D_ep_^α^ and K_ep_^α^ are shown in Figure 2. The drug uptake profiles for the base case (c_pt,ini_^α^ = 80 kg m^−3^) and the combination of D_ep_^α^ and K_ep_^α^ out of this parametric space that lead to the lowest RMSD are depicted in Figure 3. We evaluated only a discrete number of combinations of D_ep_^α^ and K_ep_^α^. As such, a more optimal combination of D_ep_^α^ and K_ep_^α^ with a lower RMSD could be found when refining even more. This step was done in section 3.2 by direct inverse modeling. However, the aim of the current section was to analyze the processes instead of finding the perfect set of parameters.

**Figure 2.**
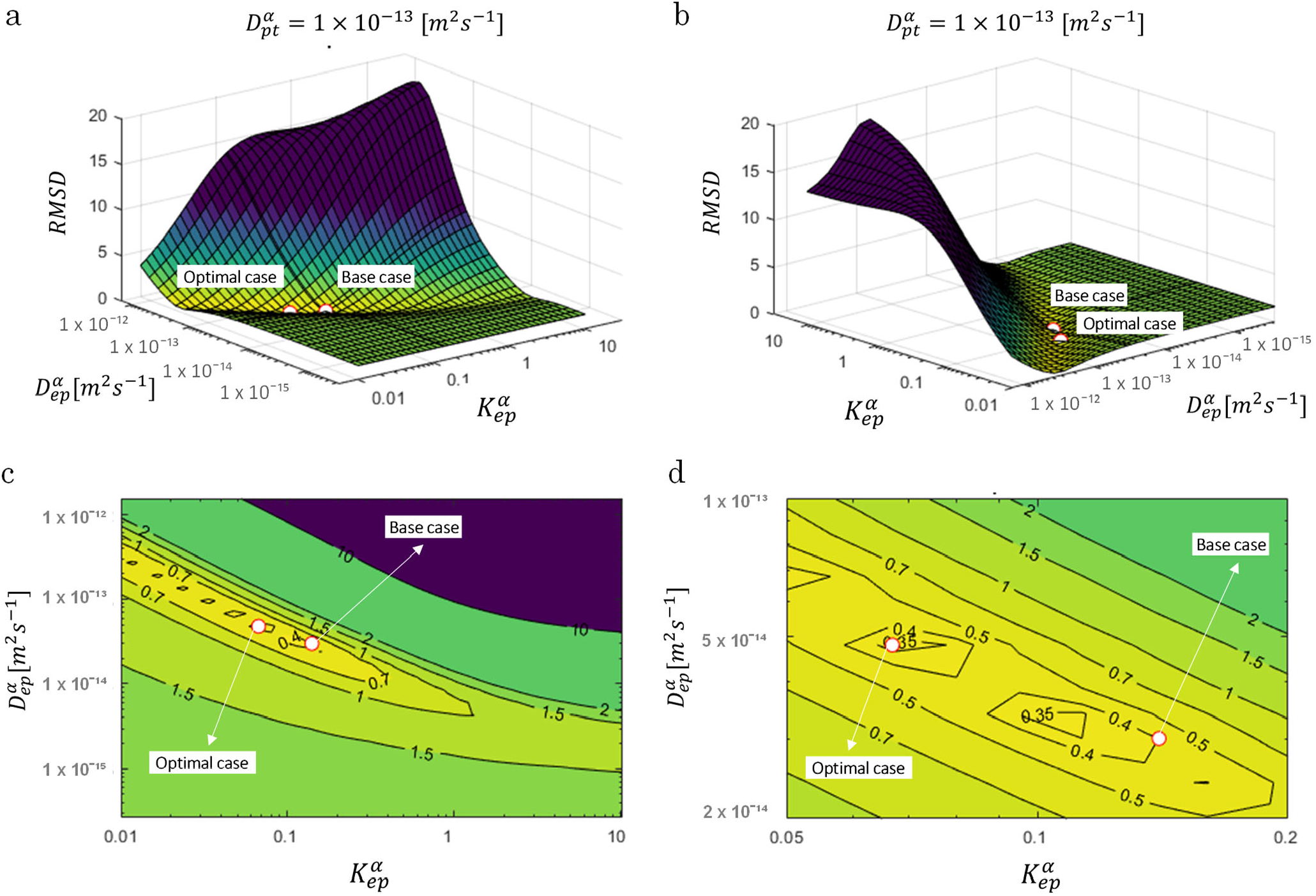
RMSD over the uptake period as a function of D_ep_^α^ and K_ep_^α^ (for D_pt_^α^ of the base case = 1.00 x 10^−13^ m^2^ s^−1^) for an initial patch concentration of 80 kg m^−3^. The parameters for the base case and most optimal parameter set are also shown.

**Figure 3.**
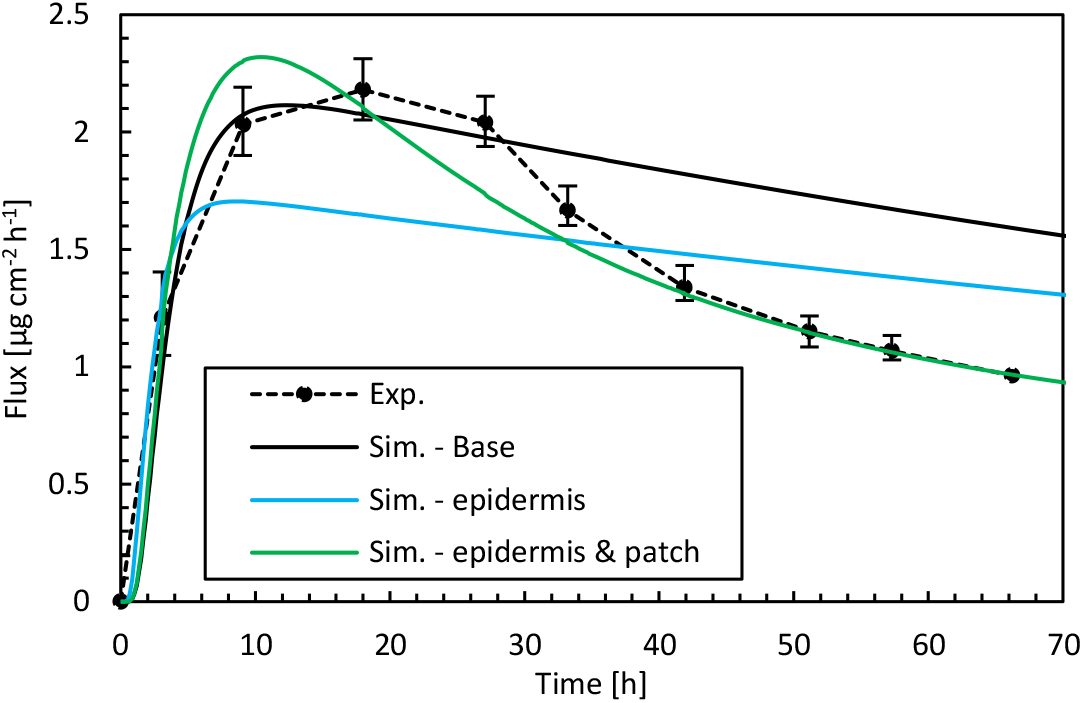
**Drug fluxes of fentanyl at surface 1** (g_bl,up_), **so leaving the epidermis, from experiments (Exp.) and simulations (Sim.) for an initial patch concentration of 80 kg m^−3^ as a function of time. Results are shown of the base case, the best combination of epidermal parameters (Sim. – epidermis, D_ep_^α^ and K_ep_^α^, for D_pt_^α^ of the base case = 1.00 x 10^−13^ m^2^ s^−1^), and the best combination of D_ep_^α^, K_ep_^α^, and D_pt_^α^ (Sim. - epidermis & patch) from indirect multiparameter inverse modeling. The fentanyl molecule is illustrated as well**.

The overall accuracy of the simulations is quantified with the RMSD (Figure 2). The RMSD of the base case (0.40) already lies quite close to the minimal RMSD value that is achieved for a more optimal combination of epidermal partition and diffusion coefficients (0.33). However, a significant improvement in RMSD was still made. The normalized RMSD was 22%, where the base case yielded 26%. In this case, the corresponding D_ep_^α^ and K_ep_^α^ were 4.8 x 10^−14^ m^2^ s^−1^ and 0.067, respectively, in contrast to 3.00 x 10^−14^ m^2^ s^−1^ and 0.14 for the base case. Interesting observations can be made when analyzing the RMSD landscape (Figure 2). A “valley/canyon” with very low values of the RMSD is present. This valley stretches over a wide range of D_ep_^α^ and K_ep_^α^. This contrasts with a typical optimization problem where an exact global minimum is present in the multiparameter space. As such, there is a broad range of possible combinations of diffusion and partition coefficients that lead to a low RMSD, so a relatively accurate simulated drug uptake process. This is problematic as an unphysical value of D_ep_^α^, for example, similar to that of the patch, can give an acceptable agreement with the experimental data when tuning/calibrating the model with the appropriate K_ep_^α^.

The simulation accuracy throughout the drug uptake process is assessed by comparing the predicted drug uptake flux over time (Figure 3). The simulations capture the initial period of the uptake accurately, namely the substantial increase in the flux. This increase occurs when drugs are transported into the epidermis. This leads to an increased drug concentration in the epidermis, so a part of the drugs is stored there. After this initial phase, drugs do not accumulate anymore in the epidermis, and a quasi-steady-state condition sets in where the amount of drugs entering the epidermis also diffuses out (Figure 4). Because the skin’s capacity to store drugs is reached, the drug primarily diffuses through the epidermis, and the stored amount remains relatively constant. Since the drug concentration in the patch (a finite reservoir) is decreasing over time, steady-state conditions with constant flux are not reached. Instead, the uptake flux slowly decreases over time, simply because the concentration gradient over the epidermis decreases. The experiments predict a much steeper decline in the flux than in the simulations, in which the predicted decline has a relatively constant slope, so is linear. During this period, the simulations do not lie within the experimental error bars. The simulations seem to miss a critical physical process, irrespective of the combination of D_ep_^α^ and K_ep_^α^ that is used. An answer is sought and found in the next section.

**Figure 4.**
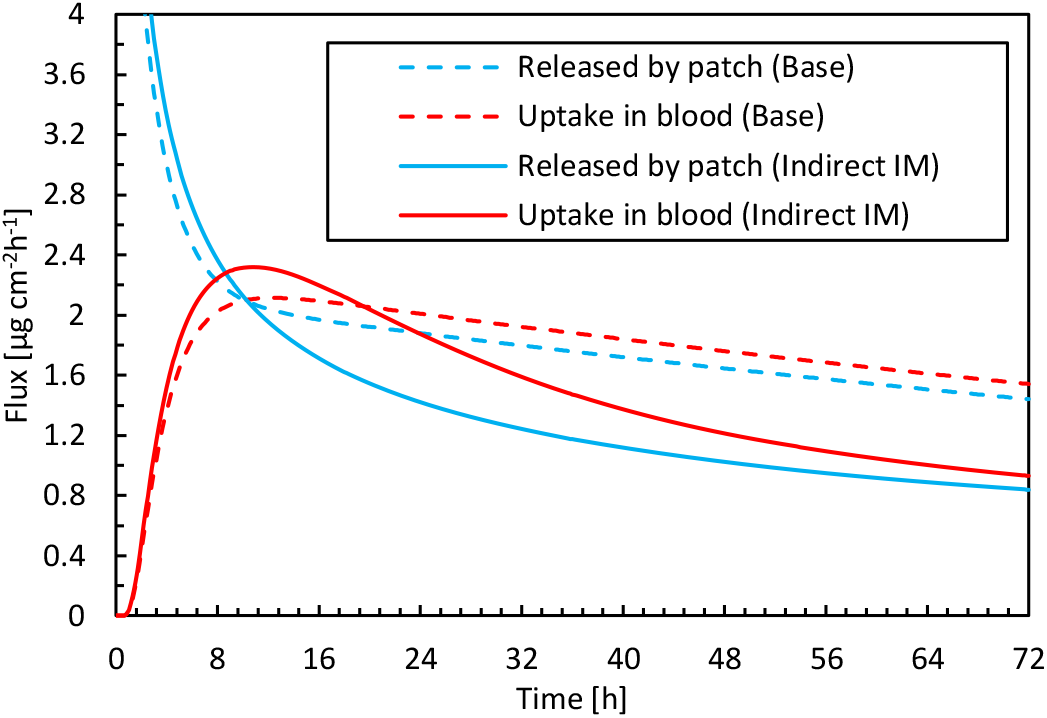
Drug flux released by the patch into the epidermis and taken up by blood as a function of time for the base case simulation and the optimal parameter combination for inverse modeling.

#### 3.1.2 The diffusion in the reservoir vs. in the epidermis

We aim to unveil whether the diffusion kinetics in the patch could help explain and improve the residual differences found between experiments and simulations in section 3.1.1. To this end, parameter variations in the diffusion coefficient of the patch D_pt_^α^ are included, in addition to those in D_ep_^α^ and K_ep_^α^. A subset of the resulting RMSD values is given in Figure 5, as depicted for different K_ep_^α^. These contour plots show how changing the ratio of the resistance to drug diffusion in the patch (D_pt_^α^), relative to that in the epidermis (D_ep_^α^), affects the drug uptake. The drug uptake profiles for the base case (c_pt,ini_^α^ = 80 kg m^−3^) and for the combination of D_ep_^α^, D_pt_^α,^ and K_ep_^α^ out of this parametric space that lead to the lowest RMSD, are depicted in Figure 3.

**Figure 5.**
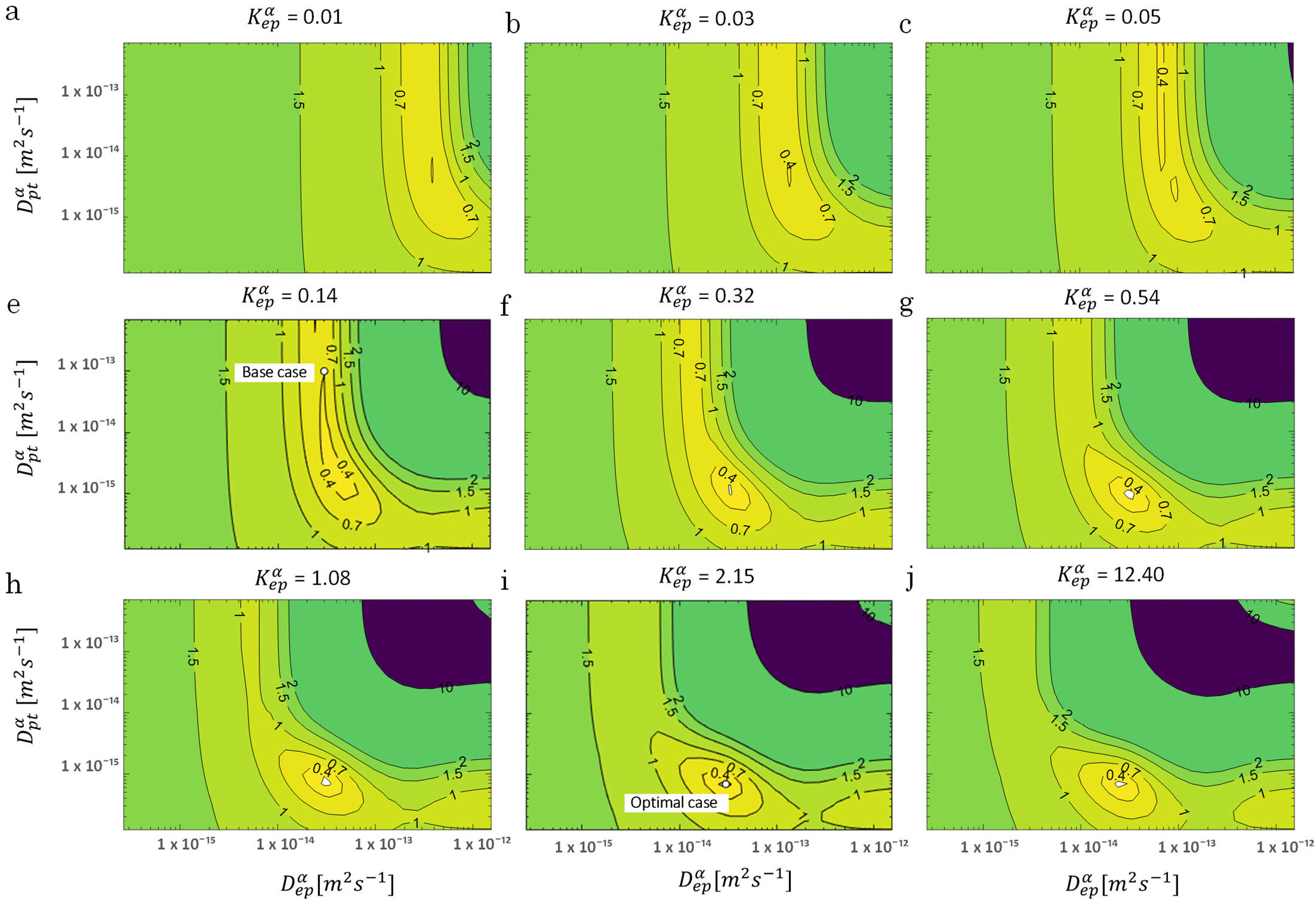
RMSD over the uptake period as a function of D_ep_^α^ and D_pt_^α^ for multiple values of K_ep_^α^ for an initial patch concentration of 80 kg m^−3^. The parameters for the base case and the most optimal parameter set are also shown.

The overall accuracy of the simulations is evaluated using the RMSD (Figure 5). A more optimal combination of D_ep_^α^, D_pt_^α^, and K_ep_^α^ made a significant improvement in RMSD (0.141) compared to only varying the epidermal model parameters (RMSD = 0.33, section 3.1.1) or the base case (RMSD = 0.4). The normalized RMSD was 9.3%. The base case yielded 26%, and the combination of D_ep_^α^ and K_ep_^α^ produced 22%. The corresponding D_ep_^α^, D_pt_^α,^ and K_ep_^α^ were in this case 3.0 x 10^−14^, 6.91 x 10^−16,^ and 2.15, respectively, in contrast to 3.00 x 10^−14^, 1 x 10^−13,^ and 0.14 for the base case. Here, “valleys/canyons” with low values of the RMSD are present in all graphs. A broad range of possible combinations of diffusion coefficients (D_ep_^α^ and D_pt_^α^) lead to a simulated drug uptake process with a low RMSD. This is true for each of the selected partition coefficients since the shape of the surface/contour plots are similar. This finding also means that partitioning between patch and epidermis does not significantly affect the relative importance of the diffusive processes in these two domains. This is confirmed in Figure 6, where the minimal values of the RMSD are shown over the full parametric space for each value of D_ep_^α^, D_pt_^α,^ and K_ep_^α^. Both D_ep_^α^ and D_pt_^α^ have an explicit minimum. However, for K_ep_^α^, there is an entire range where a RMSD below 0.15 is reached. Both values above and below one lead to a low RMSD. When the drug capacity of the epidermis is higher than one, namely the capacity of the patch (K_pt_^α^ = 1), the partition coefficient (K_pt/ep_^α^ = K_pt_^α^/K_ep_^α^= c_pt_^α^/c_ep_^α^) becomes smaller than one. This indicates that the patch’s drug concentration under equilibrium conditions is lower than the drug concentration in the epidermis. The drug will be preferably in the epidermis than in the patch, even in the absence of a concentration gradient.

**Figure 6.**
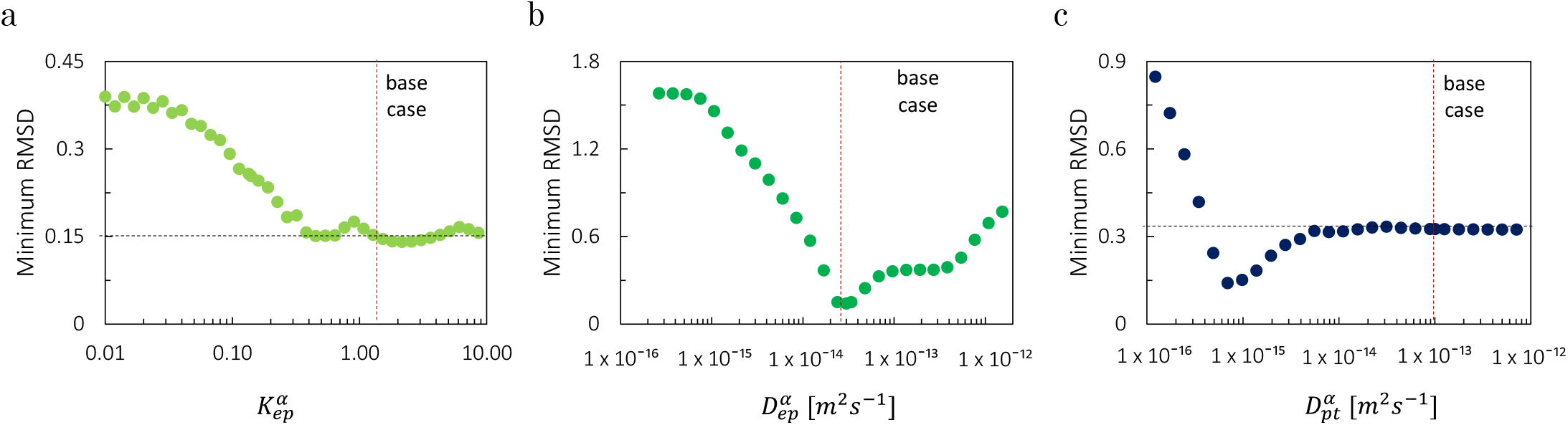
Minimal values of the RMSD over the full parametric space for each value of (a) K_ep_^α^, (b) D_ep_^α^, (c) D_pt_^α^. The values of the base case are also shown.

The simulation accuracy throughout the drug uptake process is assessed by comparing the predicted drug uptake flux over time (Figure 3). The best combination of D_ep_^α^, D_pt_^α^ and K_ep_^α^ leads to an accurate drug uptake prediction throughout the entire 72-hour timeframe, namely almost within the experimental error bars. Both the steep increase in the flux during the initial uptake period and the subsequent nonlinear decline are captured. Based on these results, we can elucidate the physical reasons for the excellent agreement for the specific set of D_ep_^α^, D_pt_^α,^ and K_ep_^α^. The lowest RMSD and the best agreement of the uptake flux profiles were obtained for a lower D_pt_^α^, compared to the base case. D_pt_^α^ was determined initially experimentally. Its magnitude resulted in much easier diffusive drug transport in the patch than in the epidermis (1.00 x 10^−13^ m^2^ s^−1^ vs. 3.00 x 10^−14^ m^2^ s^−1^). The primary resistance to diffusive transfer from the patch through the epidermis into the diffusion cell was located in the epidermis for the base case. This diffusive resistances R_i_ [s m^−1^] is defined as:

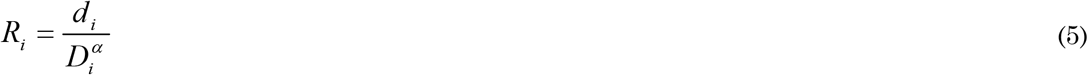

where *d_i_* is the thickness of the patch or the epidermis [m]. The corresponding resistances for patch and epidermis were 5.08 x 10^8^ and 1.69 x 10^9^ s m^−1^, respectively, for the base case. A better agreement with experiments is obtained when the diffusion in the reservoir holding the drug is restricted more so that it also has a notable resistance to drug transport. As a result of the reduced transport in the patch, even a non-uniform concentration in the patch will arise, in contrast to the base case, as depicted in Figure 7. Apart from the decreasing concentration in the patch, which leads to a reduced gradient over the skin, the concentration gradient inside the patch also plays a role. As a result, both the steep increase in the drug uptake flux and the nonlinear decline are predicted correctly (Figure 4).

**Figure 7.**
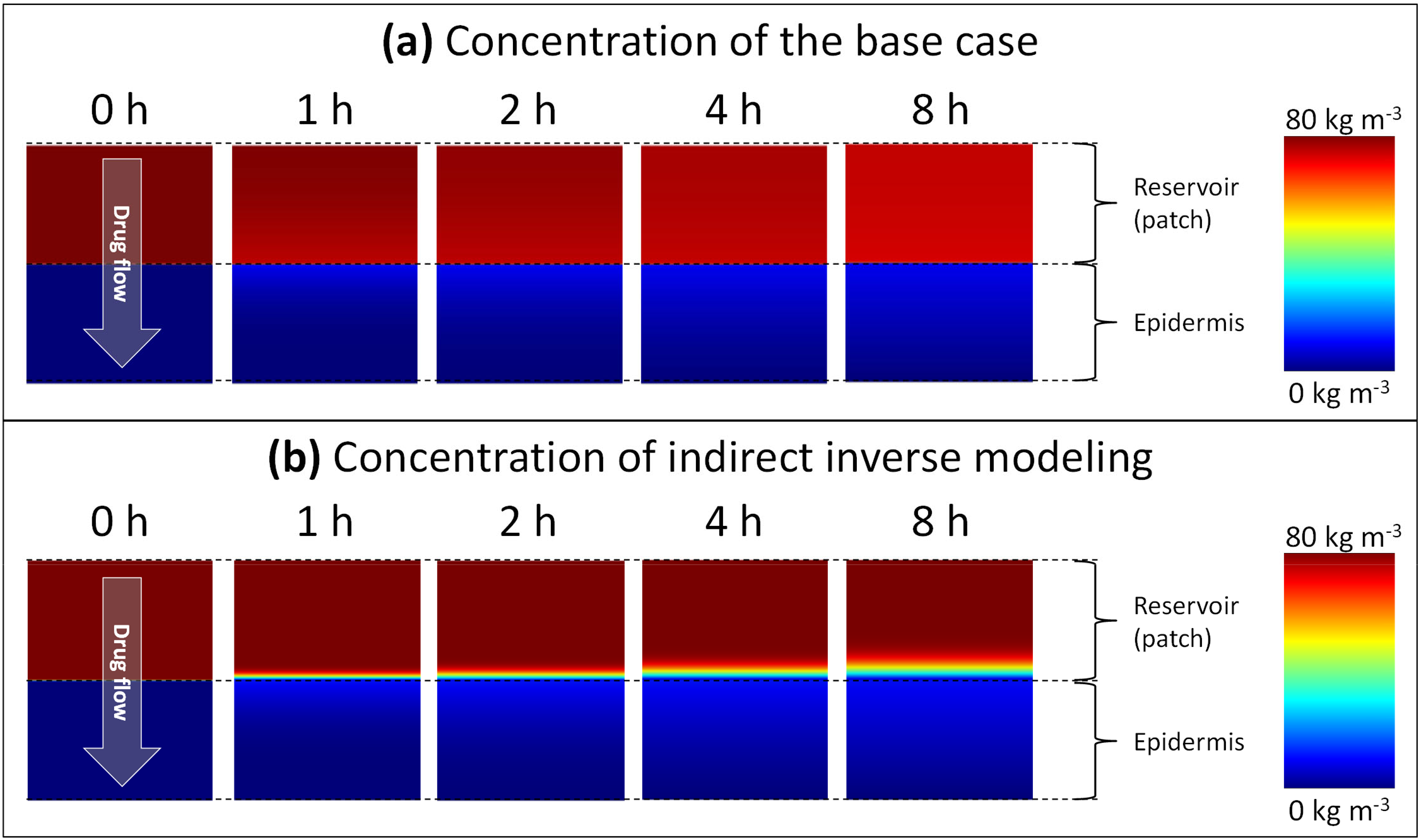
Color contours of drug concentration in the drug reservoir and epidermis for different points in time for (a) the base case and (b) simulations with optimal material properties as determined from inverse modeling.

### 3.2 Finding the optimal set of material properties by multiparameter inverse modeling

We determine the optimal set of material properties (D_pt_^α^, D_ep_^α^, K_ep_^α^) by indirect and direct multiparameter inverse modeling. For indirect modeling, we searched the combination in a sizeable parametric space of 30 618 combinations that gives the best agreement with the experimental data. The parameter set with the best agreement was identified in section 3.1.2. For direct inverse modeling, we converged in a more straightforward way to the optimal solution, without exploring the full parametric space, so with a much lower amount of simulations. The optimal combinations of material properties can be found in Table 2, together with the RMSD for each case. The drug uptake flux profile for direct and indirect inverse modeling and the RMSD at each time step are shown in Figure 8.

**Table 2.** ***Material transport properties for the base case simulations from experiments*** [18] ***and for the optimal solution for indirect and direct inverse modeling and for cross verification simulations, using the optimal parameters to simulate drug uptake with a different initial concentration***.

| Parameter | Symbol & unit | Experiment – Base case |  | Indirect inverse modeling | Direct inverse modeling |  | Cross verification |  |
| --- | --- | --- | --- | --- | --- | --- | --- | --- |
| Concentration [kg m<sup>-3</sup>] |  | 80 | 60 | 80 | 80 | 60 | 60 | 80 |
| Material properties |  |  |  |  |  |  |  |  |
| Drug capacity epidermis [-] | $K_{ep}^a$ | 0.14 | 0.14 | 2.15 | 1.59 | 0.62 | 1.59 | 0.62 |
| Diffusion coefficient epidermis [m<sup>2</sup> s<sup>-1</sup>] | $D_{ep}^a$ | $3.00 \times 10^{-14}$ | $3.00 \times 10^{-14}$ | $3.0 \times 10^{-14}$ | $2.91 \times 10^{-14}$ | $3.01 \times 10^{-14}$ | $2.91 \times 10^{-14}$ | $3.01 \times 10^{-14}$ |
| Diffusion coefficient patch [m<sup>2</sup> s<sup>-1</sup>] | $D_{pt}^a$ | $1.00 \times 10^{-13}$ | $1.00 \times 10^{-13}$ | $6.91 \times 10^{-16}$ | $7.51 \times 10^{-16}$ | $8.76 \times 10^{-16}$ | $7.51 \times 10^{-16}$ | $8.76 \times 10^{-16}$ |
| Simulation parameters |  |  |  |  |  |  |  |  |
| Number of simulations |  | 1 | 1 | 30 618 | 306 | 317 | 1 | 1 |
| Solution |  |  |  |  |  |  |  |  |
| RMSD (total) |  | 0.40 | 0.27 | 0.141 | 0.134 | 0.027 | 0.054 | 0.147 |
| Normalized RMSD |  | 26% | 24% | 9.3% | 8.8% | 2.4% | 4.8% | 10% |

**Figure 8.**
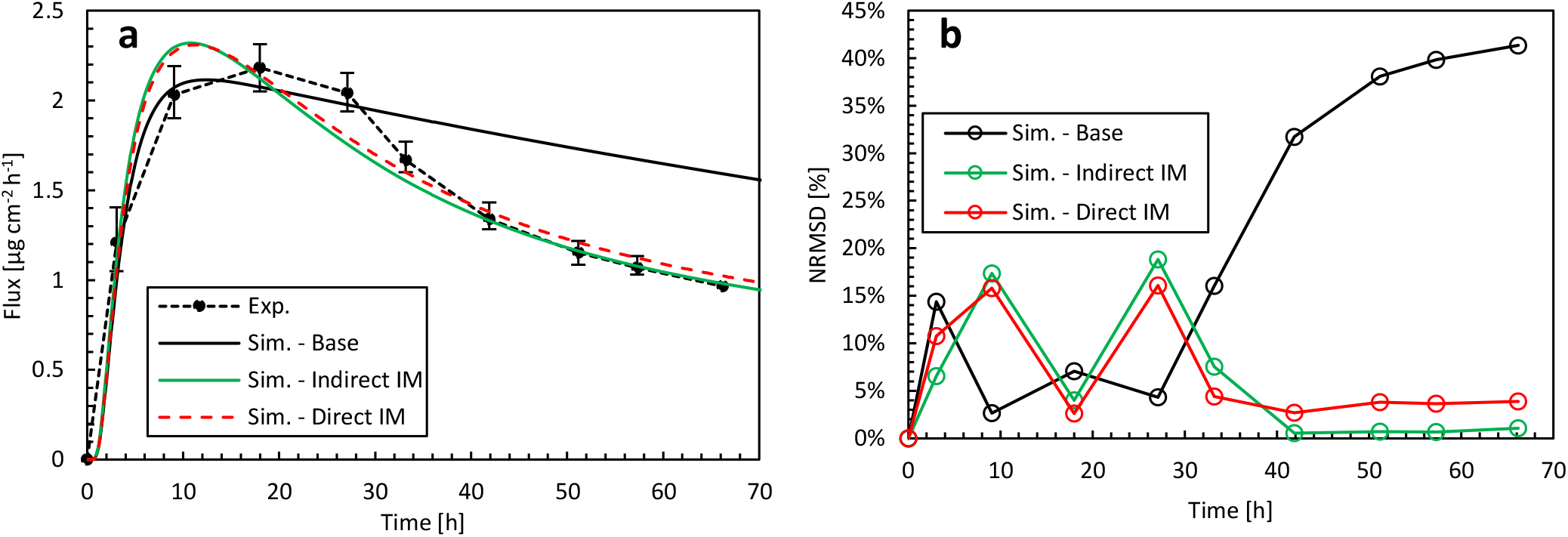
**(a) Drug fluxes of fentanyl at surface 1 (**g_bl,up_**), so leaving the epidermis, from experiments (Exp.) and simulations (Sim.) for an initial patch concentration of 80 kg m^−3^ as a function of time. Results are shown of the base case and the optimal combination of D_ep_^α^, K_ep_^α,^ and D_pt_^α^ by direct and indirect inverse modeling. (b) Corresponding RMSD at each experimental time step**.

Indirect inverse modeling gives a much better solution than with the experimentally-determined values. We improved the normalized RMSD from 26% to 9.3% and reduced the maximal deviation (per point) from 41% to 19%. Here this maximal deviation was calculated as the local NRMSD for each point (Eqs.((3),(4)), N=1). However, the sizeable parametric space is systematically explored, which is time-consuming, and only at discrete combinations of parameters D_pt_^α^, D_ep_^α^, K_ep_^α^. As only a discrete amount of combinations was analyzed, indirect inverse modeling gives the best (discrete) solution in this space but not the most optimal solution in the continuous space. This can be improved by running another parametric study around the optimal set of parameters at a higher resolution, so smaller steps between the parameter values. Nevertheless, the number of simulations to obtain this solution this way is vast.

Direct inverse modeling gives an even better RMSD than indirect modeling and much faster as well. We improved the normalized RMSD from 26% to 8.8% and reduced the maximal normalized RMSD deviation from 41% to 16%. Only 306 simulations were needed to attain the optimal solution, which is only 1% of what we had with indirect inverse modeling (30 618). Note, however, that additional simulations were performed to check the solution’s sensitivity to the starting point, which adds other computations.

Note that we also performed direct inverse modeling for c_pt,ini_^α^ = 60 kg m^−3^. The optimal combinations of material properties are also depicted in Table 2, together with the RMSD. We improved the normalized RMSD from 24% to 2.4% and reduced the maximal normalized RMSD deviation from 26% to 3.5%.

### 3.3 Cross verification of optimized model

We verify if the optimal set of transport parameters that were derived for the epidermis and patch in section 3.2 for a specific initial drug concentration (c_pt,ini_^α^ = 80 kg m^−3^), can be used for a more comprehensive operational range of patches. To this end, we simulate drug uptake with another initial concentration (c_pt,ini_^α^ = 60 kg m^−3^), for which also experimental data is available, but using the optimal parameter dataset from direct inverse modeling (Table 2), determined for (c_pt,ini_^α^ = 80 kg m^−3^). Vice versa, we also performed the same cross verification for an initial drug concentration of c_pt,ini_^α^ = 80 kg m^−3^ using the optimal model parameters obtained from direct inverse modeling for c_pt,ini_^α^ = 60 kg m^−3^. The drug uptake flux profiles for this simulation and the experimental data are shown in Figure 9, and the RMSD is shown in Table 2. The results show that there is a good agreement for both concentrations, which is in both cases better than the simulations where experimentally-determined material properties were used. For c_pt,ini_^α^ = 60 kg m^−3^ (Figure 9a), the concentration profile lies entirely within the experimental uncertainty. This means that a model, calibrated via inverse modeling on a specific concentration, can successfully be used to simulate patches with other doses as well. As such, the calibration, based on inverse modeling, only needs to be done once, and the calibrated model is valid for a more comprehensive operating range.

**Figure 9.**
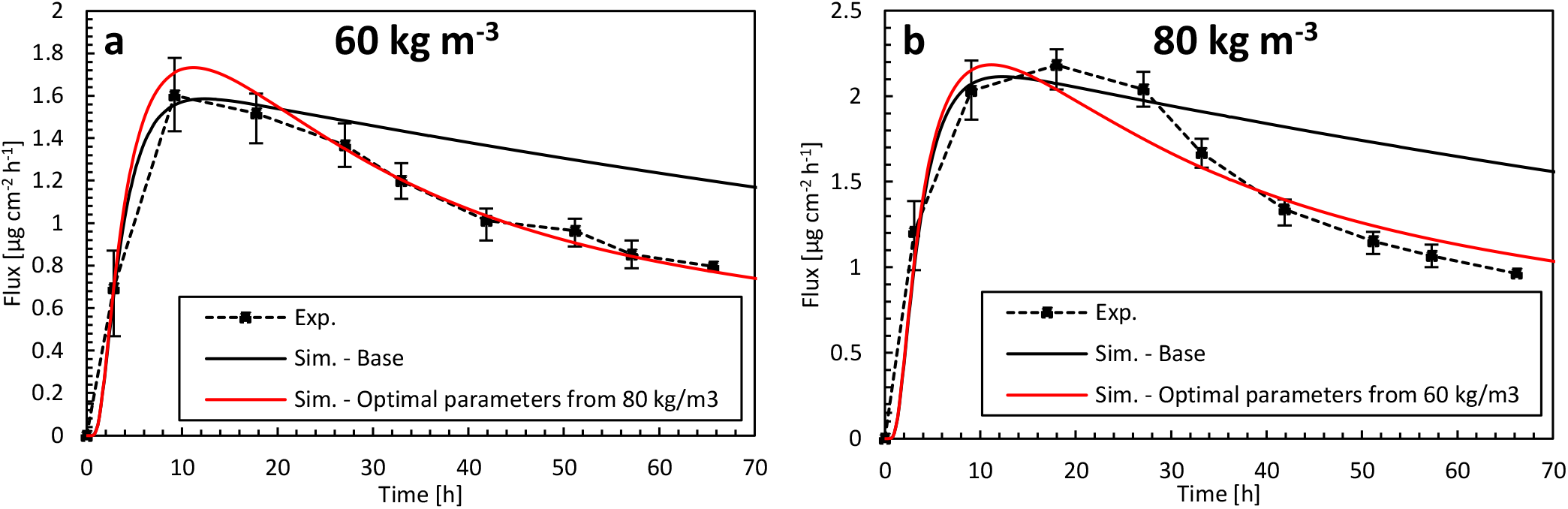
**Drug fluxes of fentanyl at surface 1 (**g_bl,up_**), so leaving the epidermis, from experiments (Exp.) for an initial patch concentration of 60 kg m^−3^ (a) and 80 kg m^−3^ (b) as a function of time and simulations (Sim.). (a) Simulation results are shown of the base case with an initial patch concentration of 60 kg m^−3^ and the optimal combination of D_ep_^α^, K_ep_^α^ and D_pt_^α^, but where the parameters are derived for an initial patch concentration of 80 kg m^−3^. (b) Simulation results are shown of the base case with an initial patch concentration of 80 kg m^−3^ and the optimal combination of D_ep_^α^, K_ep_^α^ and D_pt_^α^, but where the parameters are derived for an initial patch concentration of 60 kg m^−3^**.

## 4 DISCUSSION & OUTLOOK

A good agreement with experimental data was obtained in the present study for the optimal set of material property data, as determined by inverse modeling. This indicates that the primary physical processes at play are included in, and accurately captured by, the mechanistic model, namely diffusion and partitioning within the epidermis and transdermal patch. However, it is possible that secondary processes affect drug uptake as well, which are now not included yet. The processes that could be additionally modeled are (1) swelling or shrinkage of the epidermis caused by changing hydration due to the presence of the patch; (2) chemical metabolization (or binding) of the fentanyl molecule in the epidermis; (3) physical binding of the drug molecules so adsorption of the drug molecule into the epidermis; (4) differential diffusion in the epidermis, which can be accounted for by splitting it up into a stratum corneum and a viable epidermis, since the majority of the resistance to diffusion and the primary drug storage occurs in the SC; (5) diffusion and partition coefficients that are a function of drug concentration, instead of constant values. The importance of these secondary effects should be assessed on a case by case basis. For fentanyl, the high bioavailability through the skin (e.g., 92% in [41]) implies, for example, that physical adsorption or chemical metabolization would not play a decisive role.

The inverse modeling presented could significantly reduce experimental work and resources (typically in-vitro) that are associated with mechanistic modeling since it is no longer required to determine each of the model parameters separately. This inverse modeling only needed a single experiment on the resulting process – drug uptake via a Franz diffusion cell experiment –. An experiment at another patch drug concentration was then used for verification. Thereby, the experimental determination of the diffusion coefficients of the epidermis and patch and the partition coefficient was avoided (D_pt_^α^, D_ep_^α,^ and K_ep_^α^), which would imply three additional experiments on different equipment. The idea behind such inverse modeling to determine the material properties *in-silico* is that these properties are derived on a “training dataset” but have a sufficient accuracy to be generally applied afterward for simulations of similar cases. Verification is then used for at least one independent dataset. This can save time and resources, hence speeding up the simulation process.

The potential value of such in-silico determination of material properties becomes even larger if one wants to use mechanistic modeling for therapy for individual patients. Such mechanistic models are a fundamental building block of so-called digital twins [42]. These twins could be used for steering next-generation transdermal drug delivery systems. In clinics, significant additional resources would be required to do experiments on skin samples to calibrate the model for a specific patient. Furthermore, such lab experiments measure material properties on separate parts of the modeled system (e.g., epidermis) and different conditions (e.g., aqueous solution), which differ from reality, living human tissue. Such in-silico skin models can be used to fit the optimal model parameters, just by measuring the response of the patient to a specific biomarker that is representative of the drug of interest. Once calibrated for a particular patient, the in-silico skin model can be used to assess/design therapy within a specific operating range.

Indirect inverse modeling provided a clear insight into the impact and sensitivity of different model parameters and elucidated the optimal parameter set in-silico. Running 30 618 simulations was possible in the present study as a 1D model was used, but it will be computationally very demanding for more complex models. Direct inverse modeling is the fastest way to reveal the optimal set of parameters were and required 100 times fewer simulations to be performed. On the other hand, direct inverse modeling does not enable us to analyze and interpret the shape of the parametric space in detail, so the role of the different physics.

## 5 CONCLUSIONS

We evaluated multiparameter inverse modeling as a fast and reliable alternative to determine the drug diffusion and partition coefficients for transdermal delivery of fentanyl using mechanistic modeling. We found that capturing the drug uptake process accurately by simulations was only possible if the diffusion in the reservoir holding the drug was restricted to some extent, via D_pt_^α^, in contrast to what the experimentally-measured values suggested. This implied that a non-uniform concentration in the patch arose with distinct gradients. Only then, the initial steep increase in the drug uptake flux and the later nonlinear decline were predicted correctly.

Indirect inverse modeling was used to determine the skin and patch’s optimal material properties, namely the diffusion and partitioning coefficients. A much better agreement with the experimental drug uptake was obtained, compared to when using the experimentally-determined material properties. We improved the normalized RMSD from 26% to 9.3% and reduced its maximal deviation (per point) from 41% to 19%. A large range of appropriate combinations of diffusion and partition coefficients was identified that led to an accurately simulated drug uptake process. Hence, in this optimization problem, there was no clear minimum but rather a valley of optimal values. Direct inverse modeling gave an even better RMSD than indirect inverse modeling. We improved the normalized RMSD from 26% to 8.8% and reduced its maximal deviation from 41% to 16%. In addition, only 306 simulations were needed to attain the optimal solution compared to 30 618 with indirect inverse modeling, so a speedup of factor 100.

By cross verification, we showed that our model, when calibrated via inverse modeling for a specific concentration, can successfully be used to simulate patches with other drug doses as well. As such, the calibration, based on inverse modeling, only needs to be done once to have a valid model for a more comprehensive operating range. We thereby successfully demonstrated a fast way to build up an accurate mechanistic model for drug delivery. This opens the door to clinically-ready patient-specific therapy tailored to patients.

## Acknowledgments

This work was supported by the Novartis Research Foundation [grant “Virtual twinning for intelligent, personalized transdermal drug delivery”]. We acknowledge the support of Chandrima Shrivastava in the first exploratory work on data processing of the simulation results. The authors declare that this study received funding from Novartis Research Foundation. The funder was not involved in the study design, collection, analysis, interpretation of data, the writing of this article or the decision to submit it for publication. This manuscript has been released as a pre-print at bioRxiv.

## AUTHOR CONTRIBUTIONS

T.D. and R.R. conceptualized the study and acquired funding; T.D. did the project administration; T.D. performed the investigation, developed the methodology, performed the validation, and executed the simulations with key input from F.B.; T.D. supervised F.B.; T.D. wrote the original draft of the paper and did the visualization, with key input from F.B.; R.R. performed critical review and editing.

## Nomenclature

<u>Symbols</u>

A_pt_: active area of the patch [m^2^]
c_i_^α^: drug concentration of substance α in material i [kg m^−3^]
*c_pt,ini_^α^*: initial concentration in the patch [kg m^−3^]
d_ep_: thickness of epidermis [m]
d_pt_: thickness of patch [m]
D_i_^α^: diffusion coefficient/diffusivity of substance α in material i [m^2^ s^−1^]
g_bl,up_(t): uptake flux across the skin into the blood at a specific point in time [kg m^−2^ s^−1^]
g_Fr,up_(t): uptake flux across the skin into the Franz diffusion cell at a specific point in time [kg m^−2^ s^−1^]
K_A/B_^α^: partition coefficient between material A and B for substance α
K_o/w_^α^: partition coefficient between octanol and water for substance α
K_i_^α^: drug capacity of substance α in material i [-]
N: number of measured points
R_i_: diffusive resistance of a material [s m^−1^]
R_ep_: Radius of the epidermis in the model [m]
R_ep_: Radius of the transdermal patch [m]
S_s_^α^: volumetric source term for substance α [kg m^−3^s^−1^]
t: time [s]

<u>Greek symbols</u>

α: substance indicator
ψ^α^: drug potential of substance α [kg m^−3^]

<u>Subscripts</u>

bl: blood
i: material indicator
ini: initial
ep: epidermis
ini: initial
up: uptake
pt: patch
Sim.: In simulations
Exp.: In experiments

<u>Abbreviations</u>

TDDS: transdermal drug delivery systems
RMSD: Root mean square deviation
NRMSD: Normalized root mean square deviation

